# Assessing Bioactivity and Biointegration of Engineered Salivary Tissue Constructs in a Preclinical Unilateral Fractionated Irradiated Rat Model

**DOI:** 10.64898/2026.05.11.724009

**Authors:** Kerry P. Pernick, Juliana Amorim, Caio C. da Silva Barros, Iva Vesela, Meng-Jia Lian, Samuel Nahass, Thaise C. Geremias, Warren Swegal, Andrew M. Farach, Daniel A. Harrington, Danielle Wu, Mary C. Farach-Carson, Isabelle M.A. Lombaert

**Author notes:** **Corresponding authors:** Drs. Isabelle M.A. Lombaert and Mary C. Farach-Carson, Address: North Campus Research Center, Biointerfaces Institute, 2800 Plymouth Road, Ann Arbor, MI 48109, Address: School of Dentistry, 7500 Cambridge St. Suite 4401, Houston, TX 77054. Equal Contribution (Kerry P. Pernick, Juliana Amorim). Equal Contribution (Mary C. Farach-Carson, Isabelle M.A. Lombaert).

## Abstract

Human salivary stem/progenitor cell (hS/PC)-loaded hyaluronic acid (HA)-based hydrogels, termed 3D-salivary tissue constructs (3D-ST), hold great promise for restoring salivary gland function post-radiation injury. Here, we developed a next-generation 3D-ST using heparin-modified HA and bioactive peptide-modified hydrogels. This new formulation enables controlled pre-loading and localized presentation of heparin-binding growth factors prior to surgical implantation, providing opportunities to enhance *in vivo* hS/PC bioactivity. To model clinically relevant radiation injury, we established an athymic rat model subjected to computed tomography (CT)-guided fractionated radiation, resulting in hallmark features of radiation-induced salivary dysfunction. Over 60-days post-irradiation, glands exhibited progressive loss of acini, increased fibrosis, and disruption of endothelial, neuronal, and myoepithelial compartments. Within this injured environment, a surgical pocket was created to precisely implant 3D-STs to assess graft performance. Fluorescent labeling of the 3D-STs enabled longitudinal tracking post-implantation. Over 14 days, implanted 3D-STs remained structurally stable within irradiated glands, and hS/PCs remained viable without evidence of local inflammatory responses. Compared to non-injured glands, the irradiated microenvironment suppressed hS/PC proliferation and phenotype, indicating alterations in the irradiated local tissue negatively impact hS/PC bioactivity. In addition, host neurovascular migration into the 3D-ST was majorly restricted in irradiated glands, providing new opportunities to enhance biointegration. Overall, this work establishes a reproducible preclinical framework for assessing hydrogel biocompatibility and stability, cell bioactivity, and host-graft biointegration prior to scale up into preclinical large animal models. This study has successfully established a tractable approach for improving 3D-ST formulations to enhance hS/PC expansion, differentiation, and biointegration following implantation into radiation-injured beds.

## 1. INTRODUCTION

Implantable hydrogels are highly promising tools for tissue repair based on their mechanical properties and interactions with cells^1^. By leveraging implantation of hydrogels with encapsulation of stem/progenitor cells (e.g., of hematopoietic, mesenchymal, or epithelial lineage), one can overcome several challenges, such as low cell survival rates, poor cell engraftment, or reduced cell functionality, that often are observed in cell therapies used for a variety of diseases and injuries^2,3^.

Head and neck cancer patients receiving collateral radiation to healthy salivary glands as part of their primary cancer treatment, an event that still frequently occurs despite advances in radiation dosing and image-guided targeting, often develop hyposalivation and associated xerostomia^4^. Cell therapy is a viable reparative treatment to restore salivary gland function through repopulating epithelial saliva-producing acinar cells and their respective stem/progenitor cells^5–7^. Various hydrogels tested in the salivary gland field support these cell-related therapies. For example, hydrogels mixed with decellularized extracellular matrix can grow rat tissue bioscaffolds *ex vivo*^8^. Cell-loaded biomaterials and bioprinting approaches created human epithelial-derived organoids/spheroids/gland-like structures *in vitro*^9–18^. To date, these constructs primarily were used for *ex vivo* research purposes or for transplantation of spheroids *in vivo* after biomaterial removal^16,18,19^. Additionally, acellular biomaterials applied *in vitro* aided in murine gland repair as stand-alone gels supplemented with growth factors to promote gland repair^20–22^. However, only a few bioengineered, cell-loaded materials have been tested in the context of salivary gland regeneration *in vivo*, which include our earlier work in a ¾ resected non-irradiated rat^23^ and irradiated miniswine gland^24^. Apart from a study using an adipocyte cell-loaded hyaluronic acid (HA) biomaterial for salivary gland repair^25^, examples involving human glandular epithelial cells are limited to the implantation of organoids embedded in laminin-GelMA hydrogel^26^, collagen-encapsulated human organoids^27^, and our human salivary stem/progenitor cells (hS/PCs) encapsulated in a hyaluronic acid (HA)-based hydrogel^23,24,28–31^ (termed 3D-salivary tissue, 3D-ST) (Suppl. Table 1).

HA is a naturally occurring glycosaminoglycan abundant in human extracellular matrices and well recognized for its high hydration capacity, mechanical stability, and role in regulating cell behavior. For example, HA engages with cell-surface receptors, such as CD44 and RHAMM, participates in interactions with proteoglycans and collagenous matrices, and provides biologically relevant cues that support epithelial progenitor cell-matrix maintenance and tissue organization^32,33^. Our first-generation hydrogel used thiolated-HA (HA-SH) with a poly(ethylene glycol) (PEG) diacrylate (di-Ac, DA) (i.e., PEGDA) crosslinker to create a stable network with tunable stiffness^23^ (Suppl. Table 1). Encapsulated hS/PCs in these <u>HA-PEGDA</u> hydrogels self-assembled over time in up to 50-500 μm spheroids^30^. hS/PC spheroids expressed β_2_-adrenergic and M_3_ muscarinic receptors, and responded to neurotransmitters by increasing intracellular calcium^30^. The latter calcium fluxes are essential for proper autonomic nervous system-mediated saliva secretion and aided in the differentiation of hS/PCs into amylase-secreting acinar cells^34^.

Subsequently, we developed a second-generation peptide-modified HA hydrogel^24^ with an increased degree of thiolation to improve stability and surgical suitability. These hydrogel formulations replaced the PEGDA crosslinker with a matrix metalloproteinase (MMP)-sensitive peptide sequence^35^ (KGGG<u>PQ</u>G↓IWGQGK) reacted with Ac-PEG-succinimidyl valerate (SVA), yielding an Ac-PEG-PQ-PEG-Ac crosslinker to facilitate cell movement via cell-directed crosslink hydrolysis. The hydrogel was functionalized further with pendant Ac-PEG-SVA-modified arginine-glycine-aspartic acid (G<u>RGD</u>S) peptides^36^ (Ac-PEG-RGD) to enable integrin-mediated motility and reorganization (Suppl. Table 1). These <u>HA-RGD-PQ</u> hydrogels permit hS/PCs to dynamically remodel their local microenvironment while maintaining overall construct integrity. Using this approach, we previously demonstrated that hS/PCs encapsulated within HA-RGD-PQ hydrogels remain viable, retain amylase secretion, and grow into multicellular spheroid-like gland structures for over 8 weeks following implantation in irradiated miniswine salivary glands^24^. The latter results established the second-generation hydrogel as a biocompatible and adaptable construct for salivary tissue engineering applications.

To provide clinical translation of our 3D-ST-based cell therapy for the treatment of radiation-induced xerostomia in head and neck cancer survivors, we require durable *in vivo* hS/PC bioactivity and neurovascularization to support long-term salivary regeneration and sustain regulated secretory organ function. To enable potential priming of the 3D-STs with growth factors, we developed a third-generation hydrogel that incorporates thiol-modified heparin (HEP) within the HA-RGD-PQ network (termed <u>HA-HEP-RGD-PQ</u>). This newer formulation allows control of any pre-loading and local presentation of heparin-binding growth factors prior to and during implantation to enhance cell bioactivity or host-graft biointegration (Suppl. Table 1). To test the performance our third-generation 3D-ST in a physiologically relevant model, a cost and time effective unilateral irradiated rat model was created and validated. Using this new model, we evaluated 3D-ST biocompatibility and bioactivity after implant in healthy versus irradiated glands.

## 2. MATERIALS AND METHODS

### 2.1 Animals

Male athymic nude rats (*RH-Foxn*1*^rnu^*) were purchased from Envigo (Inotiv, West Lafayette, IN) and acclimated before any procedures were performed. The rats were 6-8 weeks old at the time of radiation and 8-10 weeks old when 3D-STs were implanted. All animal handling procedures were approved by the Institutional Animal Care and Use Committee (IACUC) at the University of Michigan. All care and housing standards, as described within the NIH Guide for Care and Use of Laboratory Animals (NIH publications No. 8023, revised 1978) were followed. A limitation of our study included the sole analysis of male rats. Even though sexual dimorphism exists in rat salivary submandibular glands^37^, no differences in parotid glands have been reported yet.

### 2.2 Fractionated Radiation Dosing Plan

A unilateral fractionated irradiation model was used, delivering 5 Gy/day for five consecutive days to the right parotid gland, for a cumulative dose of 25 Gy (X-ray exposure). Radiation was delivered using a Small Animal Radiation Research Platform (SARRP; Xstrahl Inc., Suwanee, GA, USA). Each day of irradiation, rats were anesthetized using isoflurane and safely secured vertically into the SARRP machine (Fig. 1A). A cone beam computed tomography (CBCT) image was acquired for each rat and loaded into the MuriPlan software (Xstrahl Inc., Suwanee, GA, USA) to locate the right parotid gland. An isocenter (Fig. 1A, IsoC) was placed on the identified parotid and the radiation beam was placed at an angle to avoid collateral damage to other salivary glands, the spinal cord, and vital organs in the field (Fig. 1A, blue lines). Rats were returned to housing after adequate recovery time without adverse effects. Rats were euthanized at Days 7, 14, 30, and 60 post-irradiation to allow for organ dissection and analysis for model verification.

**Figure 1.**
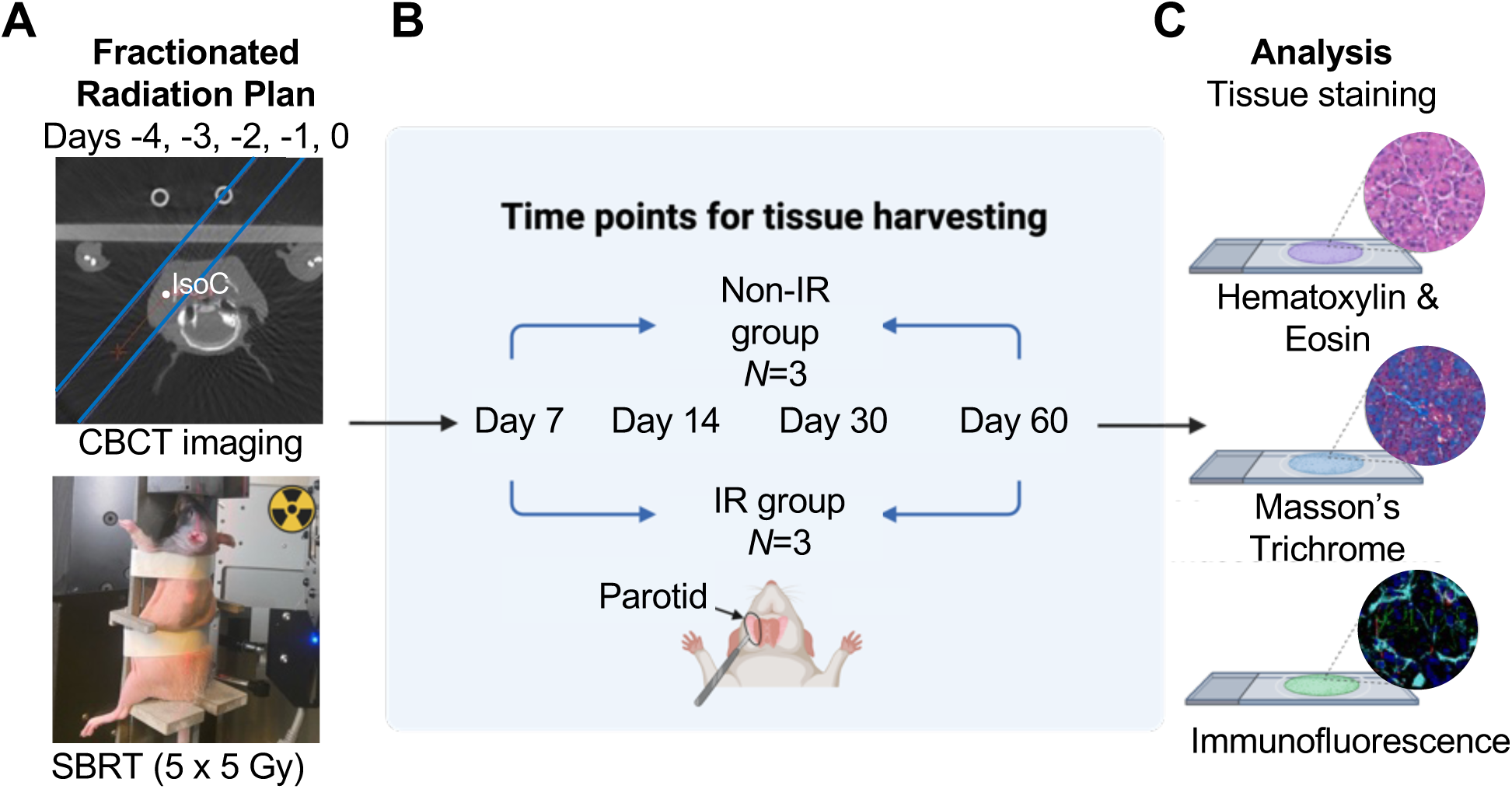
Overview of the Irradiated Athymic Rat Model. **(A)** Fractionated radiation plan highlighting a chosen radiation isocenter (IsoC) in the right parotid gland and selected beam direction (blue lines) with associated rat placement within the SBRT machine. CBCT, Cone Beam Computed Tomography. SBRT, Stereotactic Body Radiation Therapy. **(B)** Chosen time points for the collection and analysis of right parotid gland tissue from irradiated (IR) and age-matched non-irradiated (non-IR) rats. **(C)** Tissue analysis methods for verification of the animal model.

### 2.3 *Ex Vivo* Human Salivary Gland Stem/Progenitor Cell Expansion

Human parotid salivary gland specimens were collected from consenting patients (males and females, ages 22 to 81) undergoing head and neck resection at Stanford University and transferred to the UTHealth Houston under Institutional Review Board approved protocols and material transfer agreements. Freshly resected tissues were processed to isolate and expand validated hS/PCs^23,34,38^ according to a well-established protocol, as described in detail previously^28^. Briefly, tissue explants were cultured for 5-to-7 days until a 60% confluent cell sheet grew out from the explant. Cells were further expanded, passaged with trypsin, and harvested for 3D-ST hydrogel encapsulation. Low passage (< 8 passages) cells were used for experimental studies or cryopreserved. hS/PCs were maintained in culture with William’s E medium (10128-636, Quality Biological) supplemented with 1% (v/v) ITS (777ITS032, InVitria), 1 mg/mL human serum albumin (A1887, Sigma-Aldrich), 1% (v/v) GlutaMax (A1286001, Gibco), 1% (v/v) 100 U/mL penicillin and 100 µg/mL streptomycin (15140122, Gibco), 0.1 µM dexamethasone (D4902-25MG, Sigma-Aldrich), and 10 ng/mL human epidermal growth factor (hEGF) (PHG0311, Invitrogen). Media was replaced every other day.

### 2.4 3D Salivary Gland Engineered Tissue (3D-ST) Formation *Ex* Vivo

#### 2.4.1 PEGDA-Peptide Functionalization

Hydrogels were synthesized using a matrix metalloproteinase (MMP)–cleavable peptide (PQ; KGGG<u>PQ</u>G↓IWGQGK; GenScript) and an integrin-binding peptide (RGD, G<u>RGD</u>S; GenScript). Peptides were conjugated to acrylate–polyethylene glycol–succinimidyl valerate (Ac-PEG-SVA-3400, 3400 g/mol; Laysan Bio) through SVA-ester-mediated amide bond formation with primary amines to form Ac-PEG-PQ-PEG-Ac and Ac-PEG-RGD. Reactions were performed at a PEG-to-peptide molar ratio of 2:1 for PQ and 1:1.2 for RGD. The resulting PEG-peptide conjugate retained a terminal acrylate for subsequent reactions.

#### 2.4.2 Hydrogel Preparation

Thiolated hyaluronic acid (GS222F, Glycosil^®^; Advanced BioMatrix) and thiolated heparin (GS217F, Heprasil^®^; Advanced BioMatrix), were reconstituted in Buffer A (Advanced BioMatrix), incubated at 55 °C and vortexed intermittently until fully dissolved. Prior to crosslinking, the pH of each precursor solution was verified or adjusted with 1N NaOH as follows: HA 7.4, HEP 7.4, PQ 7.6, RGD 8.0. Heprasil^®^ was incorporated at 10% of the HA/HEP solution. Hydrogels were formed by combining HA-HEP, Ac-PEG-SVA-RGD, and Ac-PEG-PQ-PEG-Ac at a 4:1:1 volume ratio. The final network density was controlled by maintaining a thiol-to-acrylate molar ratio of 3:1.

#### 2.4.3 3D-ST Preparation

For *in vivo* tracking of 3D-STs post-implantation, Cy5 fluorophores (sufo-cyanine5 maleimide, 13380, Lumiprobe) were functionalized into the hydrogel with the thiol-to-maleimide molar ratio of 200:1. Cell-free hydrogels were used for initial Cy5 tracking experiments and oscillatory rheology (G’ ∼ 300 Pa). Cell-laden hydrogels were prepared with hS/PCs at 6 million cells per mL at volumes of 30-50 µL, a suitable size for the surgical pocket approach. Cell-laden 3D-STs were prepared and shipped to the University of Michigan where they were maintained in culture for 7 to 30 days prior to surgeries. All 3D-STs passed visual and tactile inspection prior to shipments to the University of Michigan and traveled well with high viability as reported previously^24^.

### 2.5 3D-ST Implantation *In Vivo*

The surgical technique used was adapted from our earlier described gland resection procedure^30^. Surgery and implantation were solely performed in the right parotid gland in all cases, regardless of whether the rat received unilateral fractioned radiation or served as (non-irradiated) control. After isoflurane-induced anesthesia, each animal was placed in a right lateral recumbent position. At the surgical site, hair was removed via chemical depilatory and skin was disinfected. A 10-15 mm incision was made using a scalpel 2-3 mm inferior to the ear and further opened using blunt surgical scissors. The layers of skin were dissected through until the parotid gland was exposed. The parotid capsule surrounding the parotid gland was partially removed to expose the gland and elevated with forceps. A shallow pocket, approximately 1-2 mm in width, was created in the gland using micro-scissors. The pocket was held open with forceps, and a 3D-ST construct was placed neatly into the pocket. The parotid gland was closed with an absorbable suture (Ethicon, Vicryl Rapide Suture, VR214) and the skin was closed with non-absorbable sutures (Ethicon, Nylon Suture, 662B). Directly after surgery and the following day, a dose of 5 mg/kg of carprofen was delivered via subcutaneous injection (VetOne, OstiFen Injection, 510510). Rats were allowed adequate recovery time and were observed carefully for 7 days post-surgery, followed by euthanasia at different time-points (up to Day 14) post-implantation to retrieve the parotid gland and implanted 3D-ST. Animals tolerated the surgical procedure well and displayed no overt signs of distress due to the implant.

### 2.6 Tissue processing, Immunostaining, Immunofluorescence, and Quantification

Harvested parotid tissue, with or without a 3D-ST, was fixed overnight in 4% (v/v) paraformaldehyde. For immunofluorescent staining, tissue was processed through an overnight sucrose gradient (% w/v: 15% sucrose overnight, 30% sucrose overnight, followed by a 1:1 ratio of 30% sucrose and Optimal Cutting Temperature (O.C.T) compound (Fisher Healthcare Tissue-Plus O.C.T. Compound, 23-730-751) overnight). The tissue was embedded in O.C.T compound and cryosectioned at 10 µm thickness. (Epredia CyroStar NX50 Cryostat, 95-726-0). Cryosectioned tissue was washed and rehydrated with 1x PBS, and a 1:1 acetone-methanol step of 5 minutes was applied. Glycine (0.1%, w/v) was applied for 15 minutes to quench residual paraformaldehyde from fixation and tissue was washed in 0.1% (v/v) Tween20/1x PBS before blocking for 2 hours in 10% (v/v) donkey serum, 1% (w/v) BSA and 0.1% Tween20/1x PBS. After blocking, primary antibodies were applied and left overnight at 4°C. The tissue was washed using 0.1% Tween20/1x PBS and secondary antibodies were applied for 40 minutes at room temperature. Tissue was washed again and mounted using Fluoro-gel (Electron Microscopy Sciences Fluoro-Gel, 17985-30). Pre-implantation 3D-ST construct staining followed a similar protocol, with Tween-20/1x PBS replaced by 0.2% TritonX-100/1x PBS, and the acetone-methanol, glycine post-treatments, and the mounting steps were omitted. Slides and 3D-ST constructs were imaged using a Nikon A1-R inverted confocal microscope. The primary antibodies used were E-cadherin (13-1900, Invitrogen), AQP5 (AQP-005, Alamone Labs), α-Smooth Muscle Actin (ACTA2) (A2547, Sigma-Aldrich), β-III Tubulin (TUBB3) (MAB1195, R&D Systems), FABP4 (AF1443, R&D Systems), Ki-67 (550609, BD Biosciences), Lamin A/C (AB133256, Abcam), Cleaved Caspase 3 (9661, Cell Signaling Technology), α-Amylase (A8273, Sigma-Aldrich), and KRT19 (Troma-III, DHSB). The secondary antibodies used were anti-mouse Cy3 (715-166-151, Jackson ImmunoResearch), anti-rat Alexa Fluor 647 (712-606-153, Jackson ImmunoResearch), anti-rabbit Alexa Fluor 488 (711-546-151 Jackson ImmunoResearch), and anti-goat Alexa Fluor 488 (705-546-147 Jackson ImmunoResearch). DAPI (D1306, Invitrogen) was used to counterstain nuclei. To quantify confocal images of immunostained slides and 3D-ST constructs, the pixel classifier tool within Qupath^39^ was used. In Qupath, annotations were drawn around each fluorescent channel to label each channel. After labeling, the pixel classifier program was used for image analysis. Each pixel in the image was labeled according to its originating fluorescent channel. The number of recorded pixels for each fluorescent channel divided by the total pixel number produced a percentage of color associated with each channel that was then used for quantitative analysis.

Paraffin embedded tissue was sectioned and stained for Hematoxylin and Eosin (H&E) and Masson’s Trichrome. A minimum of three tissue sections per stain were digitally scanned (Polaris Brightfield slide scanner). To quantify H&E and Masson’s Trichrome images, Qupath^39^ was used. To quantify the number of acinar cells, the nuclei detector tool was used to identify nuclei within selected regions of interest. Next, the object classifier tool was used to sort acinar nuclei from non-acinar nuclei (e.g., ductal cells). The number of acinar nuclei in each region of interest (ROI) was noted and used as the value for analysis. This process was repeated using nine selected ROIs on each slide. ROIs all had the same area and shape and were placed in locations with limited empty space and high density in acinar cells. To quantify the acini area, nine acini were randomly selected within each ROI, and a border was drawn around each one. The area of each acinus was recorded for each ROI within the slide and used for analysis. To quantify Masson’s Trichrome images, an ROI was drawn around each full section of tissue, excluding non-parotid tissue. The area of each ROI was recorded. To set the thresholding parameters, three colors were set as channels in Qupath to segment the image based on pixel intensity within a certain channel.

Since Masson’s Trichrome stains for three colors, these colors/channels were assigned as the blue collagen or fibrotic fibers, red muscle or cytoplasm, and dark purple/black nuclei. The number of pixels colored blue (fibrotic fibers) was recorded and divided by the area, to produce a percentage of fibrosis in the total section. This percentage then was used for further analysis.

For each analysis and to ensure a representative sampling, a minimum of three slides per glandular tissue were evaluated and three biological samples were retrieved per condition.

### 2.7 Fluorescently labeled 3D-STs to Assess Implant Stability

Before implantation, 3D-ST gels containing the Cy5 fluorophore were imaged using an IVIS fluorescence imaging machine (IVIS Lumina S5) using the appropriate emission and excitation spectrum levels (640 and 660-700 nm, respectively). On Day 7 post-implantation, rats were placed under the IVIS imaging machine with the implantation incision still intact. Images were taken using the same settings as the gel pre-implantation. Next, the external skin incision was opened, and additional IVIS images were taken with the internal parotid suture intact or after glandular suture removal.

### 2.8 Statistical Analysis

Quantitative data collected from histological scoring and immunostaining were analyzed using GraphPad Prism 10.6.1. The Shapiro-Wilk test was used to test all data for normality. Welch’s unpaired Student’s *t*-test was used for normal data, and the Mann-Whitney test was used for non-parametric data. For comparisons involving multiple time points, independent comparisons were performed between (non-irradiated) control and irradiated groups at each time point, as appropriate. Statistical significance is indicated on the graphs (see asterisks positioned above corresponding comparisons). Significance was set at *P* < 0.05.

### 2.9 Illustrations and Diagrams

Schematic figures and diagrams were crafted with BioRender.com using a licensed account.

## 3. RESULTS

### 3.1 Establishment of the Clinically Relevant Irradiated Athymic Rat Model

With no other examples of a clinically relevant irradiated athymic rat model available, a CT-guided unilateral fractionated dosing plan was developed for athymic rats by specifically targeting the right salivary parotid gland. During the process, collateral radiation damage to other vital organs in the CT field was carefully monitored and avoided (Fig. 1A). To generate a clinically relevant radiation dosing plan with injury outcomes, the right parotid gland was dosed for 5 constitutive days with a fractionated dose of 5 Gy. Right parotid gland tissue was removed at Days 7, 14, 30, and 60 from euthanized irradiated animals for subsequent injury analysis and compared to age-matched healthy non-irradiated animals (Fig. 1B-C). Hematoxylin & Eosin (H&E) and Masson’s Trichrome staining were performed to evaluate the level of acinar cell loss, acini area, and initiation of fibrosis (Fig. 2A-C). At all measured time points, non-irradiated rat parotid glands demonstrated numerous lobular parenchyma that each consisted of densely packed serous acini around ducts, supported by thin, visibly healthy, connective stroma in between gland lobules and within lobes (Fig. 2A, Suppl. Fig. 1A-C). By 7 Days after the last day of fractionated dosing, multiple acini appeared smaller and showed atrophic alterations (Fig. 2A, Suppl. Fig. 1D). Despite no significant reductions observed in acinar cell number at this time point as compared to healthy glands, acini area was reduced significantly by Day 7 (Fig. 2C). Similarly, parotid acini showed profound vacuolization by Day 60, and inter-acinar spaces became visually abundant over time (Suppl. Fig. 1D). The number of acinar cells started to significantly decrease after Day 14, as compared to healthy non-irradiated glands (Fig. 2C). This steady decline ultimately resulted in a significant acinar cell loss of 12% within 60 days (as compared to Day 7). A similar decline in acini area was noted wherein more than 50% of their surface area was lost by 60 days post-radiation (Fig. 2C). Accumulation of fibrosis significantly increased from Day 7 through Day 60 and visually presented itself strongly in between remaining ducts and acini as well as profoundly in between lobules (Fig 2B, blue, Suppl. Fig. 1E, arrows). H&E staining further revealed a variable but progressive increase in chronic mononuclear inflammation over time, characterized by mild inflammatory cell infiltration within the tissue at Day 7 that became increasingly prominent by Days 14 and 30, with widespread inflammation within the parotid tissue and blood vessels by Day 60 (Suppl. Fig. 1F, arrows).

**Figure 2.**
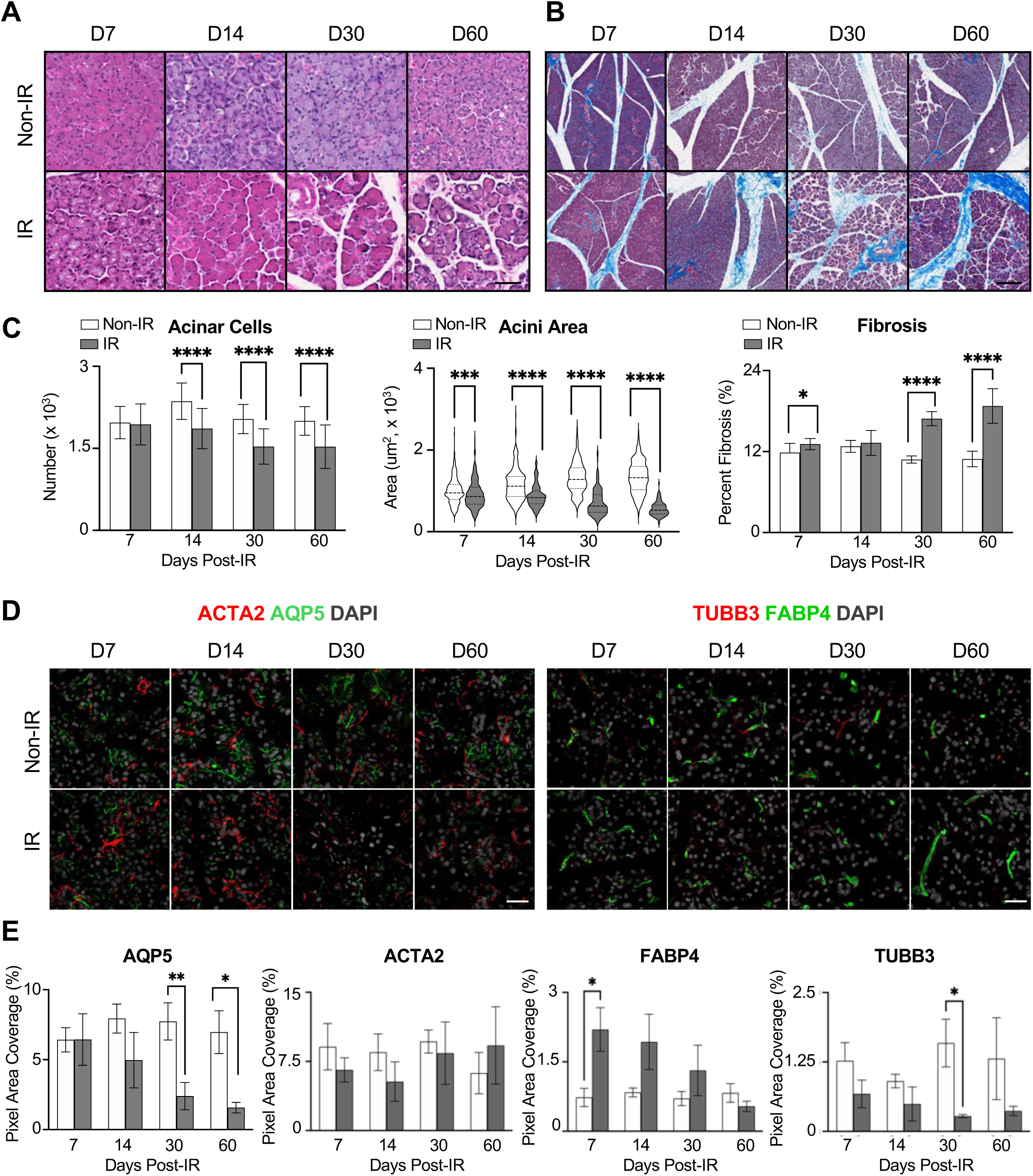
Analysis of the Clinically Relevant Fractionated Radiation-induced Rat Model. **(A)** H&E staining on right parotid glands of irradiated (IR) and age-matched non-irradiated (non-IR) rats at each collected timepoint. Scale bar, 50 µm. **(B)** Masson’s Trichrome staining on right parotid glands of non-IR and IR rats at each timepoint. Scale bar, 200 µm. **(C)** Quantification of acinar cell number, acini area, and level of fibrosis detected in tissues from A-B. Mean ± SD. *, *P* < 0.05, ***, *P* < 0.001, ****, *P* < 0.0001. **(D)** Right parotid gland from non-IR and IR animals at each timepoint immunostained for α-smooth muscle actin (ACTA2, red), Aquaporin 5 (AQP5, green), Fatty Acid Binding Protein 4 (FABP4, green), β-III Tubulin (TUBB3, red), and DAPI (grey). Scale bars, 20 µm. **(E)** Quantification of total acinar cell-related AQP5 protein, myoepithelial cell-related ACTA2 protein, endothelial cell-related FABP4 protein, and neuronal cell-related TUBB3 protein expression. Mean ± SD. *, *P* < 0.05. **, *P* < 0.01.

To assess functional changes within specific irradiated parotid gland cell types, Aquaporin-5 (AQP5, acinar cells), α-smooth muscle actin (ACTA2, myoepithelial cells), β-III-tubulin (TUBB3, neuronal cells), and Fatty acid binding protein 4 (FABP4, endothelial cells) immunostaining was performed. A significant decrease in AQP5 protein was observed from Day 30 onward, with AQP5 staining exhibiting an altered morphology marked by a reduction in its characteristic distribution on the apical membrane (Fig. 2D-E, Suppl. Fig. 1G, arrow). In contrast, a rapid and significant increase in ACTA2, related to myoepithelial cells in the epithelial compartment (Suppl. Fig. 1H, arrow), was noted at Day 7 post-radiation that subsequently subsidized to control levels again by Day 60 post-radiation (Fig. 2D-E). No statistically significant change could be observed in endothelial-related FABP4 protein across all time points (Fig. 2D-E). However, similar to our previous findings in irradiated mouse submandibular glands^40^, morphological changes in blood vessels were observed across irradiated time points, including a significant increase in blood vessel length by Day 60 post-IR (Suppl. Fig. 1I, arrows) and reduction in blood vessel number by Day 14 (Suppl. Fig. 1J). On the other hand, TUBB3 protein expression did steadily decrease over time, indicating a loss in innervation (Fig. 2D-E, Suppl. Fig. 1I, asterisks).

Given these collective findings, a Day 14 post-irradiated parotid bed was chosen for all subsequent 3D-ST implantation experiments because of its clinically relevant outcomes with a significant loss in acinar cells, innervation, and morphological changes in blood vessels, but before major fibrosis integrated the tissue.

### 3.2 Advancing *In Vivo* HA-PEG Implantation in Athymic Irradiated Glands

To conduct 3D-ST implantation, a neck incision was made to expose the right parotid gland in athymic rats, as done previously^23^ (Fig. 3A). To accommodate the 3D-ST implant, a pocket was created in the parotid tissue bed of irradiated or (non-irradiated) control rats (Fig. 3B-C). This pocket procedure proved more reliable than our previously used method wherein the 3D-ST was placed on top of the exposed parotid^23^. After the 3D-ST was transferred into the pocket under sterile conditions, a follow up suture in the parotid gland with additional skin sutures closed the incision and held the 3D-ST firmly in place (Fig. 3D-E). The surgical procedure was reproducible and well tolerated in the athymic rat model, allowing for normal healing within one week after surgery and no sign of infection (Fig. 3F). Even though the irradiated parotid gland showed a trend in decreasing weight due to gland atrophy (Suppl. Fig. 2), sufficient tissue remained such that the pocket 3D-ST implantation technique was still feasible. Unexpectedly, a variable decreasing trend in parotid gland weight also was observed in non-irradiated athymic rats over the 60-day period, even though glands were visibly healthy. This finding indicates that assessing tissue damage, rather than measuring gland weight, is a better index of radiation sequelae when characterizing salivary glands scheduled to receive implants.

**Figure 3.**
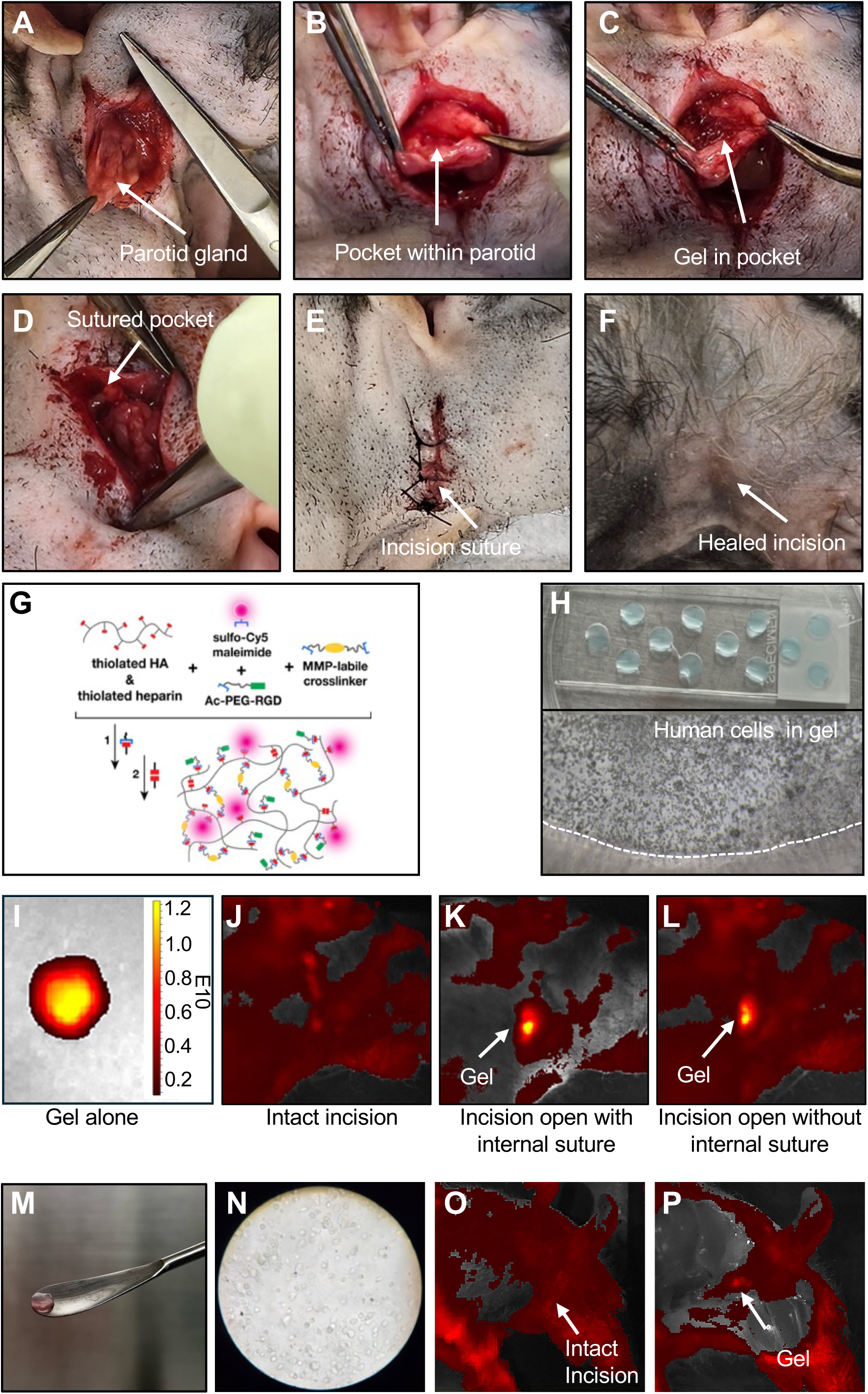
Surgical Implantation and Live Stability Testing of 3D-ST into the Irradiated Parotid Bed. **(A)** Skin incision to expose the right-side parotid gland. **(B)** Creation of a small pocket into the gland. **(C)** 3D-ST implantation into the glandular pocket. **(D)** The pocket is sutured to enclose the 3D-ST. **(E)** The surgical site is closed by additional sutures. **(F)** Representative image of healed incision 7 days after implantation. **(G-H)** 3D-ST fabrication schematic: molding of HA-HEP-RGD-PQ constructs, hS/PCs after encapsulation in hydrogel, and final 3D-ST assembly. **(I)** Fluorescent imaging of a representative Cy5-labeled hydrogel with hS/PCs before implantation. **(J-L)** Fluorescent imaging of implanted Cy5-labeled hydrogel in intact animal (J), after opening the incision site (K), and after removing the internal parotid suture (L) at 7 days post-implantation. **(M)** 3D-ST construct with hS/PCs without Cy5 labeling. Scale bar, 50 mm. **(N)** Presence of hS/PC-derived spheroids in the Cy5-labeled 3D-ST, 10 days after *in vitro* assembly and before implantation. **(O-P)** Fluorescent imaging of animal with Cy5-labeled 3D-ST before opening the incision site (O), and after opening the incision site (P) at 7 days post-implantation.

In summary, we generated a reproducible 3D-ST implantation technique for use in either the non-irradiated or irradiated parotid bed of athymic rats to allow robust assessment of *in vivo* biomaterial stability and hS/PC bioactivity.

### 3.3 Assessing Biomaterial Stability in the Irradiated Gland

To allow in-live tracing of the implanted biomaterial, a Cy5 fluorophore was added to 3D-ST hydrogels during the crosslinking process (Fig. 3G), which provided structurally consistent constructs with a light blue appearance (Fig. 3H). Cy5-labeled hydrogels, without hS/PCs, produced a high and consistent fluorescent output (∼ 1.2 x 10^10^ arbitrary units) (Fig. 3I) before implantation into a 14-Day post-irradiation parotid bed. Seven days post-implantation, a low signal was observed from the incision site but no *in vivo* fluorescent output from the implanted hydrogel could be detected (Fig. 3J). The loss of Cy5-hydrogel signal has been observed before by other groups when attempting to label hydrogels with commercially available fluorophores^41^, and has been attributed to *in vivo* instability of fluorophores by biological enzymes or degradation of the hydrogel itself. Nevertheless, when the suture site was re-opened to expose the Cy5-hydrogel, it was stably embedded and intact in the sutured irradiated parotid tissue bed (Fig. 3K). Upon removal of the glandular suture, the Cy5-hydrogel remained clearly visible with Cy5 detection and illustrated the poor skin penetrance of the Cy5 fluorophore (Fig. 3L).

We next sought to determine if Cy5 affected hS/PC behavior in 3D-STs shown at the time of implant (Fig. 3A). When comparing regular 3D-STs with or without Cy5-labeled 3D-STs (Fig. 3H and 3M), the formation of hS/PC spheroids *ex vivo* was visually not influenced by the integration of Cy5 into the hydrogel (Fig. 3N). As expected, the presence of the hS/PCs in the hydrogel did not alter quenching of the Cy5 signal by the skin after glandular implantation, and a positive Cy5 emission from the hydrogel remained after opening the skin incision site at 7 days post-implantation (Fig. 3O-P).

These data indicate that the presence of Cy5 in the 3D-ST allows for rapid detection of the exposed cell-loaded hydrogel and permits observance of the gradual hydrogel degradation within the extracted irradiated tissue, especially when associated glandular sutures absorb. However, the Cy5-hydrogel is less suitable for IVIS imaging without skin removal. Nevertheless, no movement of the implant in the pocket was observed at any study termination, allowing easy identification of the hydrogel in the parotid bed for further hS/PC bioactivity and biointegration analysis.

### 3.4 Determining Bioactivity and Biointegration of 3D-STs *In Vivo*

3D-STs embedded in parotid glands were isolated at various timepoints and processed for immunostaining (Fig. 4A). At 14 days post-implantation, viable hS/PC nuclei were readily identified within the 3D-ST, which formed a clear 3D integrated perimeter with the irradiated rat tissue (Fig. 4B). Morphologically, no immediate inflammation was observed around the 3D-ST perimeter, indicating stable tissue integration of the implanted construct. Human-specific Lamin A/C protein staining confirmed the presence and human nature of hS/PCs within the gland-embedded 3D-ST (Fig. 4C-D). Biocompatibility of 3D-STs at Day 14 post-implantation was assessed in non-irradiated and irradiated rat parotid glands through proliferative marker, Ki-67, and early apoptosis marker, cleaved Caspase 3 (CASP3). A significant reduction in proliferating Ki-67^+^ hS/PCs was observed in a downwards trend from pre-implantation to post-implantation in (non-irradiated) control glands to irradiated glands (Fig. 4E-F, Suppl. Fig. 3A). Positive levels of CASP3 were identified in hS/PCs of 3D-STs after implantation in both healthy and irradiated glands, but no significant differences were observed between both post-implantation conditions. However, in pre-implantation conditions, significantly less CASP3^+^ hS/PCs were observed in 3D-STs. The latter observation indicates that after 3D-ST implantation, CASP3 protein is activated in hS/PCs despite cells retaining nuclei with non-apoptotic features (Fig. 4E-F, Suppl. Fig. 3A). To evaluate whether the phenotype of hS/PCs is retained post-implantation, 3D-STs were stained for ductal marker Keratin19 (KRT19) and acinar secretory protein Amylase1 (AMY1). Pre-implantation, hS/PCs retained their original undifferentiated (KRT19^-^) ductal phenotype^34^ or spontaneously differentiated into Amylase1 (AMY1^+^) acinar cells. Interestingly, a mixed population of both KRT19^+^ ductal and AMY1^+^ acinar cells were formed in 3D-STs implanted in non-irradiated parotid beds, which reflected a healthy-like morphology where KRT19^+^ ducts were surrounded by well-organized AMY1^+^ acini (Fig. 4G, arrow). Despite both KRT19^+^ ductal cells and AMY1^+^ acini being formed in 3D-STs implanted in irradiated beds, their morphological structure was less organized and a decreased trend towards AMY1^+^ acinar formation was observed (Fig. 4G-H). Host biocompatibility of the 3D-ST before and after gland implantation was assessed using expression of neuronal marker β-III Tubulin (TUBB3) and blood vessel marker Fatty acid-binding protein 4 (FABP4). *In vitro*, hS/PCs lacked expression of FABP4 but occasionally expressed low levels of TUBB3, an observation previously described in mitotic human skin keratinocyte and fibroblasts during cell culture^42^. Rat-derived neurovasculature was scarcely present in the 3D-ST, and no statistical differences were seen between the parotid bed conditions (Fig. 4I-J). Interestingly, a strong presence of neuronal fibers was detected at the 3D-ST border with the non-irradiated parotid bed (Fig. 4K). However, the denervating irradiated parotid bed showcased a decreased trend of neuronal fibers at the 3D-ST border (Fig. 4L). In comparison, rat-derived blood vessels were abundantly present at the 3D-ST border, independent of the parotid bed it was inserted into (i.e., irradiated and non-irradiated) (Fig. 4L).

**Figure 4.**
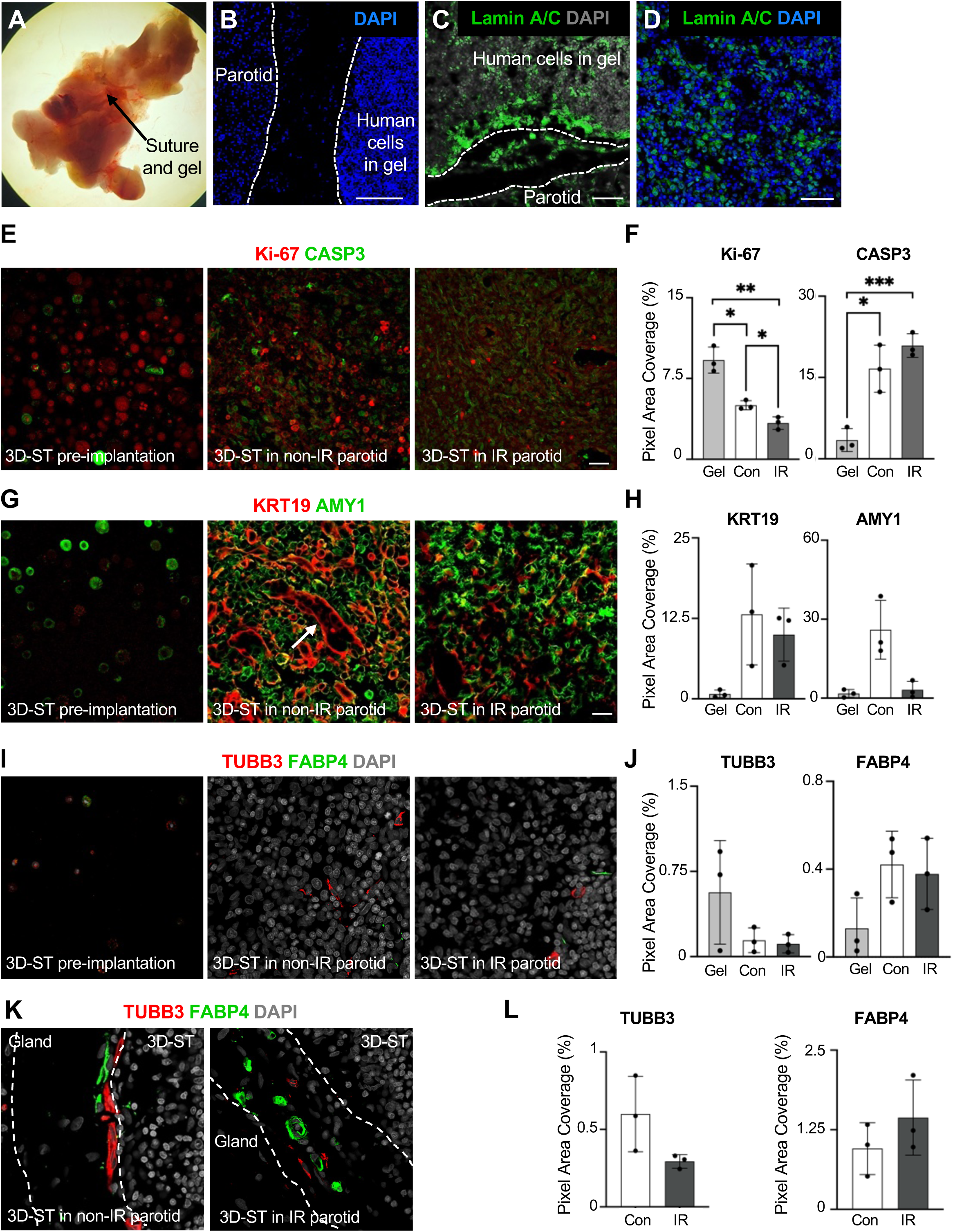
*In vivo* Compatibility of 3D-STs Post-implantation. **(A)** Representative image of an irradiated parotid gland with implanted Cy5-labeled 3D-ST and glandular suture to hold the 3D-ST in place at Day 14 post-implantation. **(B)** Nuclear staining (DAPI, blue) of extracted 3D-ST with surrounding rat irradiated parotid gland (from A). Scale bars, 100 µm. **(C-D)** Immunostaining of 3D-ST and surrounding rat irradiated parotid gland (from B) for human-only Lamin A/C (green) and DAPI (blue). Scale bars, 20 and 50 µm. **(E)** Immunostaining of a 14 Day implanted 3D-ST for Caspase 3 (CASP3, green) and Ki-67 (red) in a 3D-ST construct pre-implantation, or after implantation in a non-irradiated (non-IR, Control, Con) and irradiated (IR) parotid bed. Scale bar, 20 µm. **(F)** Quantification of Ki-67 and CASP3 protein staining in (E). **(G-H)** Keratin19 (KRT19) and Amylase1 (AMY1) protein expression and quantification in pre-implanted and 14-day post-implanted 3D-STs in a non-irradiated and irradiated parotid bed. Scale bar, 10 µm. **(I-J)** Staining and quantification of neuronal (TUBB3) and endothelial (FABP4) protein expression in pre-implanted and 14-day post-implanted 3D-STs in non-irradiated and irradiated parotid beds. Scale bar, 10 µm. **(K-L)** Visualization and quantification of TUBB3^+^ neuronal fibers and FABP4^+^ endothelia at the 3D-ST border with the non-irradiated or irradiated parotid bed at 14 days post-implantation. Mean ± SD. *, *P* < 0.05. **, *P* < 0.01 (for F, H, J and K).

In summary, hS/PCs remain viable in the visibly non-immune provoking HA-HEP-RGD-PQ hydrogel. However, a significant reduction in hS/PC proliferation and ductal-to-acinar-ratio morphology was observed as compared to 3D-STs embedded into healthy glands, illustrating the hydrogel effectively diffuses inhibitory signals, or lack thereof, from the irradiated tissue bedding to hS/PCs. Additionally, host biointegration into the 3D-ST was limited but substantial neurovasculature could be detected at the 3D-ST border.

## 4. DISCUSSION

In this study, we first developed a robust method to assess the *in vivo* stability and biocompatibility of HA-HEP-RGD-PQ hydrogels with or without fluorescent labels for implantation into radiation injured parotid beds. This successful approach further allowed us to initiate analysis of hS/PC bioactivity and host-graft biointegration. Immunocompromised and immunosuppressed animals are essential components in preclinical *in vivo* studies to determine the regenerative success of human cell-based biomaterials, but very few of the existing models have assessed biointegration in radiation-damaged tissues. Here, we created a new clinically relevant athymic rat salivary gland injury model using a unilateral fractionated radiation scheme.

Histologically, major salivary glands consist of 80% saliva secretory acini^43^, which are connected to a robust ductal structure that allows for transportation of saliva into the oral cavity. For optimal salivary functionality, both acini and ducts must be properly surrounded by a basement membrane and interact with proximal myoepithelial cells, along with connections to endothelial, neuronal, and stromal cells in the associated interstitial compartment^44^. However, radiation therapy has a disproportionate and immediate negative impact on acinar cells that are highly radiation-sensitive and is often associated with an increase in fibrosis and reduction in the neuronal and blood vessel compartment at emerging chronic stages^45–47^. Over time, irradiated immunocompetent rat parotid acinar cells typically show progressive vacuolization with inter-acinar spaces due to cell loss^48^ and dilated ducts^49^. Acinar abnormality and functionality often are associated with a downregulation of AQP5, which is needed for water secretion, in the apical membrane of acinar cells^50^.

In a similar way, our data reveals that a unilateral fractionated radiation plan to the parotid glands of athymic rats rapidly results in morphological changes in the irradiated tissue. The significant acinar cell loss, increase in fibrosis, and decreasing trend in innervation reflect both acute and chronic responses, which has been widely seen in the clinic^45–47^. Nevertheless, in our model, we observed both acute and chronic injury characteristics occurring in a shorter timeframe of 60 days (2 months), as compared to humans where chronic stages initiate at 3 months and continue later. Thus, our irradiated athymic rodent model rapidly and accurately reproduces the most important challenges to restoring function to the radiation-damaged parotid and is, hence, an excellent model for evaluating the regenerative success of 3D-STs.

To design translationally feasible repair strategies for irradiated glands, we adapted the implantation technique from our previous gland resection model for use in athymic rats suitable for implantation of human hS/PCs. In our earlier work, ¾ of healthy parotid glands were resected and 3D-STs were embedded immediately adjacent to the remaining ¼ residual parotid bed with subsequent wound closure^23^. In this study, we developed a reliable ‘pocket’ method to determine the *in vivo* stability and biocompatibility of implanted 3D-STs. The surgical pocket approach secured the hydrogel containing hS/PCs in a position where it lies directly within the parotid, and is readily subjected to the host’s neurovascularization network. By integrating a Cy5 fluorochrome into the hydrogel formulations, 3D-STs could be readily visualized post-implantation, which demonstrated stable integration into the parotid bed without signs of inflammation, regardless of the condition of the parotid. Unexpectedly, the skin barrier and incision site quenched much of the fluorophore, precluding use of fluorescent imaging in live animals to follow implant stability, but rapid hydrogel detection in the 3D embedded irradiated gland at the time of gland/gel resection was possible. Our results indicated that implanted hS/PCs behaved robustly in Cy5-labeled hydrogels as in non-labeled cell-loaded constructs.

The use of various hydrogels, extracellular matrix components and fibers has been applied previously to initiate repair in radiation-injured salivary glands^9–18^, which has significant clinical value for head and neck cancer survivors suffering from xerostomia. Use of hydrogels in lab-based studies improved the secretory phenotype of human salivary gland-derived epithelial cells by increasing 3D organization, acinar differentiation, and response to neurotransmitters in formed spheroids, as compared to 2D monolayers^16,34^. Similarly, transplantation of human 3D spheroids, used after *ex vivo* biomaterial removal, into irradiated glands showed enhanced engraftment, when compared to their single cell counterparts^16^, and initiated reduced apoptosis in neighboring cells in the surrounding irradiated tissue bed. The latter indicates that *ex vivo* created 3D human gland-derived spheroids provide better therapeutic potential than single cells do, both *in vitro* as well as after *in vivo* transplantion^16,18^. However, not all single cells or cells within 3D spheroids survive after transplantation in the irradiated tissue bed^16^. Thus, encapsulating cells in supportive biomaterials can provide additional cell protection and positively influence human stem cell self-renewal and differentiation capacity.

In this sense, the choice of hydrogel matters. For example, it was reported that 3D collagen hydrogels induce human salivary gland-derived epithelial duct cells to form a lumen and enhance acinar-related mucin and amylase production over that of cells encapsulated into Matrigel^®27^. In contrast, human gland epithelial spheroids created on PEG-nanofibrous microwells maintained a stem cell-like property without measurable differentiation before implantation^18^. Likewise, human gland-derived cells that formed organoids in laminin-GelMA hydrogels remained undifferentiated unless dexamethasone, Bone Morphogenetic Protein (BMP), and N-[(3,5-Difluorophenyl)acetyl]-L-alanyl-2-phenylglycine-1,1-dimethylethyl ester (DAPT) was added to the media^26^. Our original HA-PEGDA hydrogels allowed epithelial hS/PCs to differentiate into amylase-producing acinar cells after treatment with neurotransmitters and supported acinar polarity that is needed for proper apical-based saliva secretion^30^. However, these formulations did not support assembly into higher order branched structures, leaving our second-generation iterations to incorporate peptides containing RGD-PQ motifs for cell migration^24^ and now a third-generation HA-HEP-RGD-PQ for incorporation of stromal-activating growth factors to enhance cell bioactivity and host biointegration.

While a number of human salivary gland-derived organoids and cell-loaded hydrogels have been used for in lab research applications (reviewed in^3^), few have been translated to *in vivo* studies that analyzed cell bioactivity and host biointegration. Transplantation of human gland spheroids encapsulated in collagen matrices into irradiated (7.5 Gy) immunodeficient mice showed engraftment into the irradiated tissue bed of salivary submandibular glands up to 4-16 weeks post-transplantation while maintaining expression of ductal and acinar markers and lumen formation^27^. In comparison, our 3D-STs similarly showed successful engraftment in healthy^23^ and irradiated parotid beds of rats and miniswine^24^ from 2-to-8 weeks. Post-implantation, no immune rejection was seen, hS/PCs remained viable, and spheroids were visibly growing. After 3D-ST implantation into the irradiated athymic rat parotid bed, hS/PCs also remained viable in the non-immune provoking HA-HEP-PQ-RGD hydrogel. However, a significant reduction in hS/PC proliferation was observed as compared to 3D-STs embedded into healthy glands and pre-implantation 3D-STs. It is interesting to observe that 3D-ST implanted hS/PCs expressed a detectable level of cleaved Caspase-3, a protein typically associated with early apoptosis. Emerging evidence revealed Caspase-3 can play non-apoptotic roles in various cell types, including stem/progenitor cells, to facilitate tissue regeneration through cell survival, proliferation, and differentiation^51,52^. Further research will be needed to reveal the precise role Caspase-3 plays in hS/PCs pre- and post-implantation. While hS/PCs have the potential to differentiate into KRT19^+^ ductal and Amylase1-producing acinar cells, the morphological formation of ducts with surrounding acini mimicked that of healthy-like glands when transplanted into a healthy gland. However, that same morphological acinar-to-duct connection was not maintained in 3D-STs embedded in irradiated glands, suggesting that the irradiated bed currently limits proper formation of new glandular constructs. Other studies revealed that the host environment dictates the bioactivity of transplanted cells^53^. In addition, neurovascular integration from the host gland into the 3D-ST was found to be scarce. Yet, the fact that the host neurovasculature is abundant nearby the 3D-ST border with the irradiated and non-irradiated parotid bed indicates that current cues to attract the neurovasculature into the 3D-ST are too low or absent. Thus, upcoming studies will need to focus on optimizing the biointegration of 3D-STs and hS/PC bioactivity in the rat irradiated bed in the short- and long-term as a step toward efficient testing in large animal models^24^.

## 5. CONCLUSIONS

This study established a reliable rat model of parotid unilateral fractionated radiation wherein we demonstrated the stable integration and compatibility of 3D-STs in parotid beds. The inclusion of a Cy5 fluorophore into the cell-loaded biomaterial enabled rapid localization of the 3D-ST into the irradiated bed. In addition, we observed that the irradiated bed negatively influenced the survival, bioactivity, and biointegration of the 3D-ST, warranting steps for future testing of growth factor-containing hydrogels.

## CREDIT AUTHORSHIP CONTRIBUTION STATEMENT

**Kerry Pernick**: Investigation, Formal Analysis, Data Curation, Writing, Visualization. **Juliana Amorim**: Investigation, Visualization. **Caio da Silva Barros**: Investigation. **Iva Vesela**: Investigation. **Meng-Jia Lian**: Investigation. **Samuel Nahass**: Investigation. **Thaise Geremias**: Investigation. **Danielle Wu**: Methodology, Review & Editing. **Daniel Harrington**: Methodology, Review & Editing. **Warren Swegal**: Methodology, Review & Editing. **Mary C Farach-Carson**: Conceptualization, Methodology, Resources, Review & Editing, Funding Acquisition. **Isabelle Lombaert**: Conceptualization, Methodology, Validation, Resources, Writing, Visualization, Review & Editing, Funding Acquisition.

## ACKNOWLEDGEMENTS

This work was supported by the National Institutes of Health (R01 DE032364, MPI: Farach-Carson M. and Lombaert I.). We thank Dr. Quynh-Thu Le’s team for providing human biopsies (Standford Medicine, CA, USA). We thank Dr. Dave Karnak for assistance with the SBRT machine (University of Michigan, MI, USA).

## STATEMENT OF SIGNIFICANCE

The novelty of our work lies in the evaluation of biostability, bioactivity, and biointegration of hydrogel-containing human epithelial stem/progenitors transplanted in healthy versus irradiated parotid gland beds. This is significant for upcoming translational work of this cell-containing hydrogel for the repair of irradiated glands in patients suffering from xerostomia.

## SUPPLEMENTARY MATERIALS

**SUPPLEMENTAL FIGURE 1, related to FIGURE 2. (A)** Region of interest (ROI) selected within a scanned rat parotid gland section. Arrow, inter-glandular stroma. Scale bar, 200 µm. **(B)** Outline of acinar and non-acinar cells within the ROI. Individual areas within the ROI were selected to measure acini. Scale bar, 50 µm. **(C)** Outline of a duct, measured acini, and intra-glandular stroma. Scale bar, 10 µm. **(D)** Higher magnification of a Day 7 and Day 60 post-IR rat parotid gland, indicating atrophied acini and inter-acinar spaces (yellow arrows). Scale bar, 25 µm. **(E)** Higher magnification of Day 60 non-IR, Day 14 and Day 60 post-IR rat parotid glands stained with Masson’s Trichrome to outline the accumulation of fibrosis in between various epithelial cell types (yellow arrows). White asterisk, intra-glandular fibrosis. Scale bar, 50 µm. **(F)** Higher magnification of H&E outlining the infiltration of immune cells in parotid glands across time (yellow arrows). Scale bar, 50 µm. **(G)** Immunostaining of Day 7 and Day 14 IR parotid glands for ACTA2 (red), AQP5 (green), and DAPI (grey). Arrows, apical location of AQP5 on acini. Scale bar, 20 µm. **(H)** Immunostaining of Day 7 IR parotid glands for E-cadherin (green) and ACTA2 (red) to outline ACTA2-expressing cells are epithelial-derived. Scale bar, 10 µm. **(I)** Staining of TUBB3 (red), FABP4 (green), and DAPI (gray) in Day 7 control and Day 60 IR parotid glands. Arrows, differences in blood vessel morphology. White asterisk, presence of TUBB3^+^ neuronal fibers. Scale bar, 20 μm. **(J)** Quantification of the number of FABP4^+^ blood vessels in parotid glands of non-IR and IR mice at various timepoints. Mean ± SD. *, *P* < 0.05.

**SUPPLEMENTAL FIGURE 2, related to FIGURE 3**. Analysis of the right parotid gland weight in non-IR and IR animals at various time-points, normalized to body weight. Mean ± SD.

**SUPPLEMENTAL FIGURE 3, related to FIGURE 4. (A)** Immunostaining for Caspase 3 (CASP3, green) and Ki-67 (red) in a 3D-ST construct pre-implantation, or after implantation in a non-irradiated and irradiated parotid bed with DAPI. Scale bar, 20 µm. Similar images as in Figure 4E. **(B)** Keratin19 (KRT19) and Amylase1 (AMY1) protein expression and quantification in pre-implanted and post-implanted 3D-STs in a non-irradiated and irradiated parotid bed, counterstained with DAPI. Scale bar, 10 µm. Similar images as in Figure 4G.

**SUPPLEMENTAL TABLE 1.** Table outlining the various generations of hydrogels referenced in this manuscript, each with their own characteristics.

## DISCLOSURE ON ARTIFICIAL INTELLIGENCE TOOLS

During the preparation of the work, the authors used QuPath pixel, cell, and object classification in order to quantify and analyze data. After using this tool/service, the authors reviewed and edited the content as needed and take full responsibility for the content of the publication.

