## Supplemental Table and Figures for "Assessing Bioactivity and Biointegration of Engineered Salivary Tissue Constructs in a Preclinical Unilateral Fractionated Irradiated Rat Model"

**SUPPLEMENTAL TABLE 1**

| Hydrogel Used in 3D-ST Transplantations | Gel Formulation | Features Identified | References |
|---|---|---|---|
| **HA-PEGDA**<br>(HA-SH & Ac-PEG-Ac) | • Thiolated HA (HA-SH)<br>• Crosslinked with poly(ethylene glycol) di-acrylate (Ac-PEG-Ac or PEGDA) (Glycosil, BioTime, ESI BIO). | 1) Biocompatible and stable in resected healthy athymic rat parotid<br>2) Tunable stiffness<br>3) hS/PCs form spinning spheroids that form lumens<br>5) Spheroids deposit basement membrane at periphery<br>6) Spheroids express CD44 and RHAMM<br>7) G' range ~60–300 Pa | Pradhan-Bhatt *et al.*, 2014<br>https://pubmed.ncbi.nlm.nih.gov/23832678/<br><br>Wu *et al.*, 2019<br>https://pubmed.ncbi.nlm.nih.gov/31750298/ |
| **HA-RGD-PQ**<br>(HA-SH$^{high}$ & Ac-PEG-PQ-PEG-Ac & Ac-PEG-RGD | • Higher thiolation levels in HA-SH (Glycosil, ESI BIO) with varying thiol–acrylate molar ratios (SH:Ac, 6:1, 3:1) to control crosslinking densities.<br>• Crosslinked with PEGDA and proline-glutamine (PQ) peptide sequence to form a modified metalloproteinase (MMP)-degradable substrate for cells. Peptide GGG**PQ**↓IWGQGK, GenScript) was conjugated to PEG (Ac-PEG-SVA, 3.4kDa, LysanBio).<br>• Additional functionalization of arginine-glycine-aspartic acid (G**RGD**S, RGD) peptides to Ac-PEG-succinimidyl valerate (Ac-PEG-SVA) (3.4kDa, LysanBio). | 1) Biocompatible and stable in immunocompromised miniswine parotid<br>2) Tunable stiffness<br>3) Slowly degradable<br>4) hS/PCs migrate within gel<br>5) hS/PCs form spheroids and irregular structures<br>6) G' range ~300 Pa | Wu *et al.,* 2021<br>https://pubmed.ncbi.nlm.nih.gov/34660692/ |
| **HA-HEP-RGD-PQ**<br>(HA-SH$^{high}$ & Ac-PEG-PQ-PEG-Ac & Ac-PEG-SVA-RGD & Heprasil) | • HA-RGD-PQ with 10% Heprasil®-Advanced BioMatrix (i.e., thiolated heparin) to control pre-loading of heparin-binding growth factors. | 1) Biocompatible and stable in athymic rat healthy and irradiated parotid<br>2) Tunable stiffness<br>3) Slowly degradable<br>4) hS/PCs migrate within gel<br>5) hS/PCs form spheroids and irregular structures<br>6) Can be labeled with Cy 5 for ready visualization<br>7) Can be readily loaded with heparin-binding growth factors<br>8) G' range ~300 Pa | Pernick *et al.* (this manuscript) |

#### SUPPLEMENTAL FIGURE 1, related to FIGURE 2

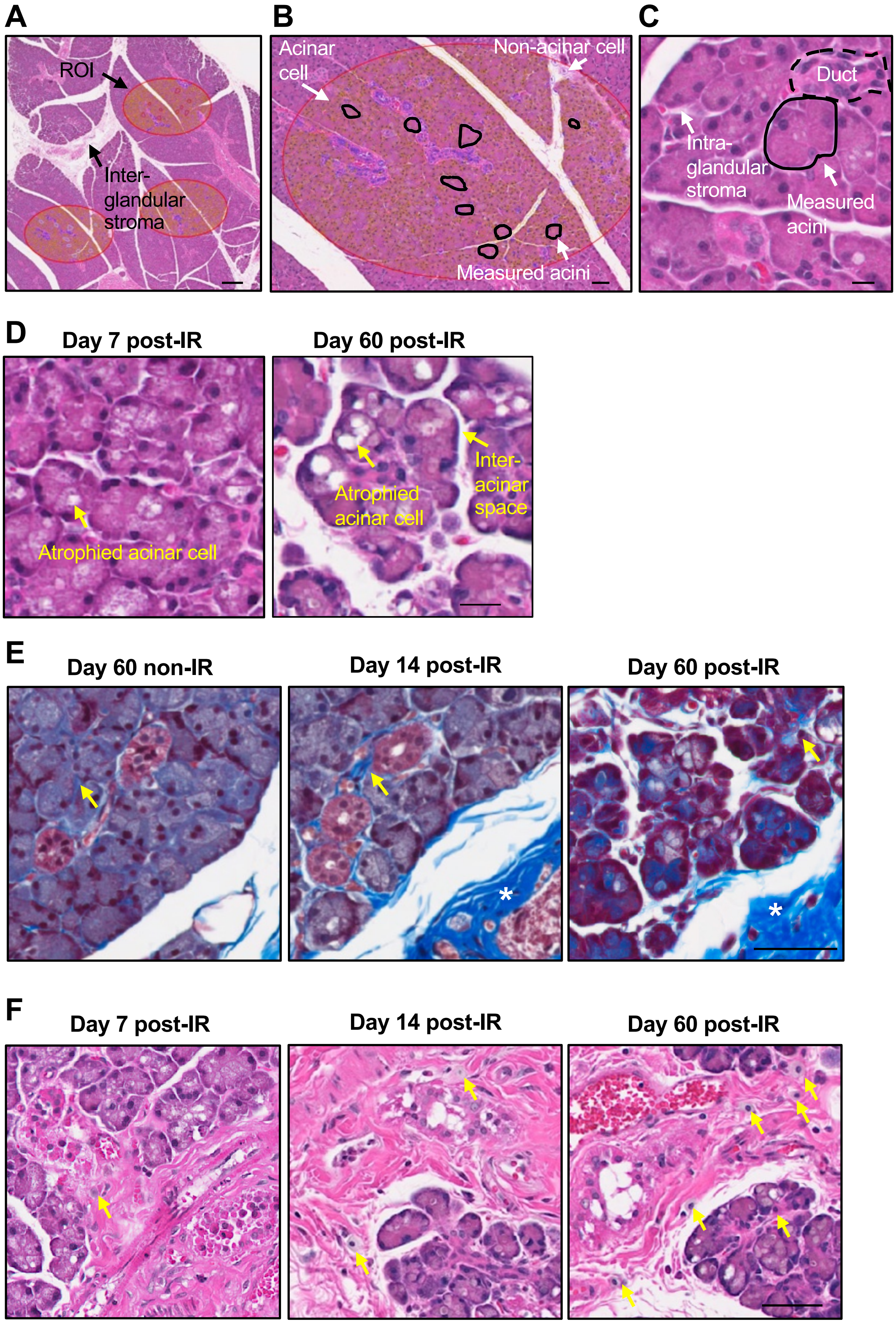

### SUPPLEMENTAL FIGURE 1, related to FIGURE 2

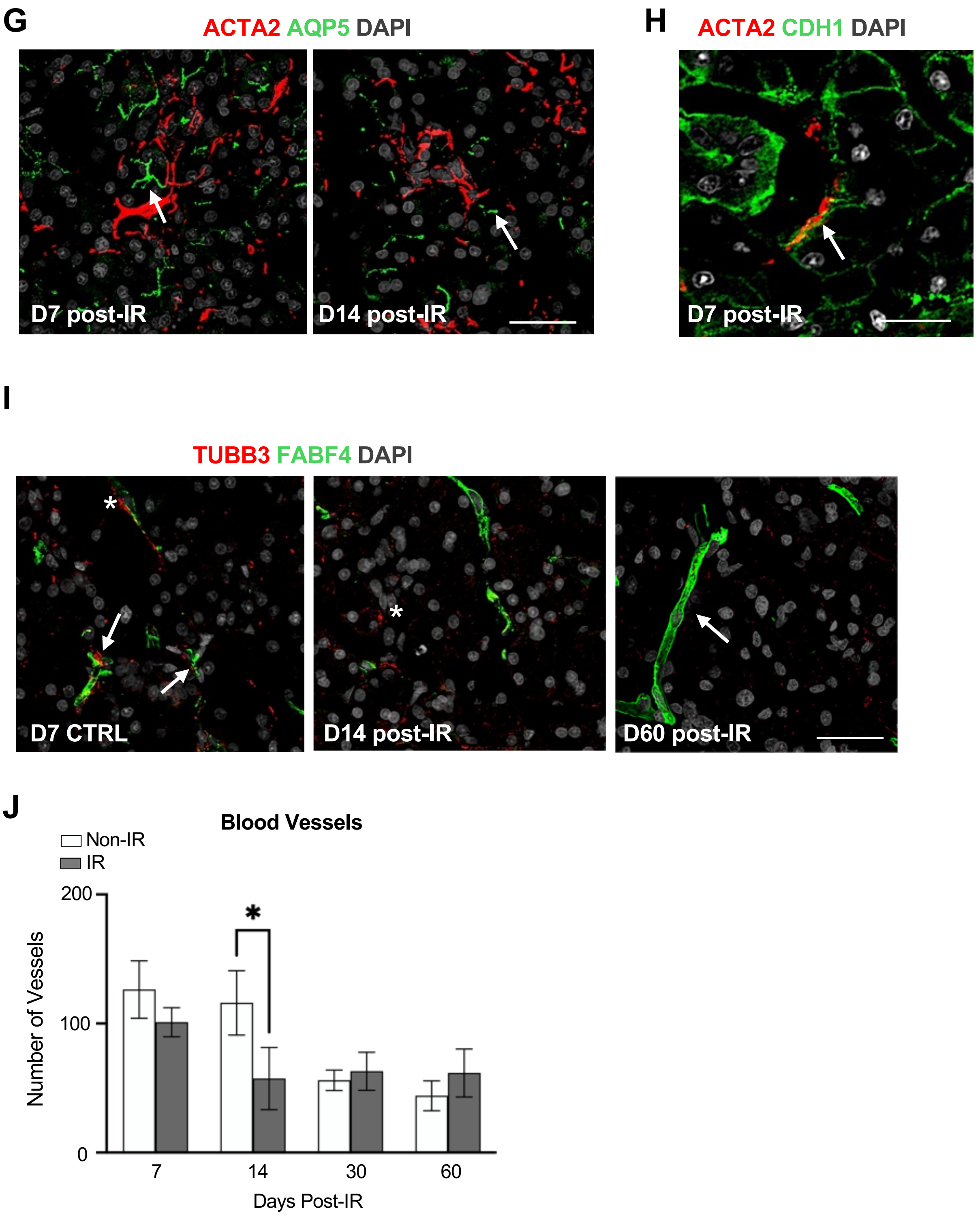

### SUPPLEMENTAL FIGURE 2, related to FIGURE 3

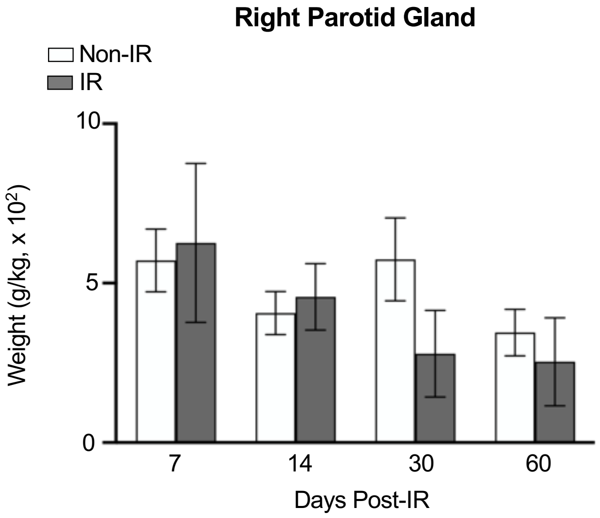

**SUPPLEMENTAL FIGURE 3, related to FIGURE 4**

A

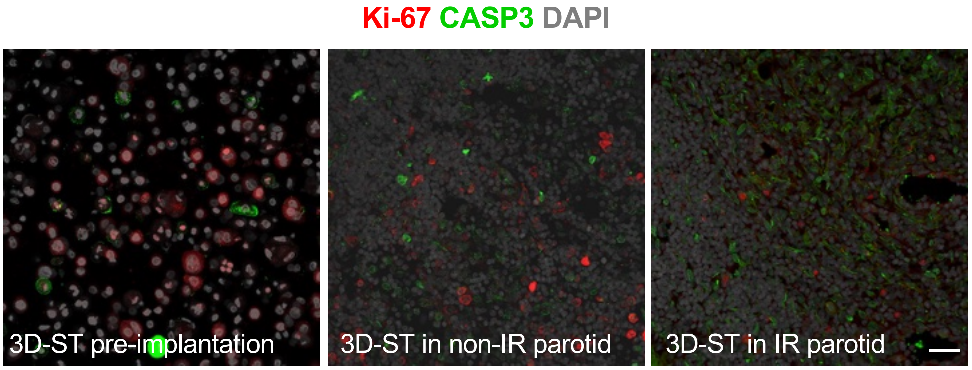


B

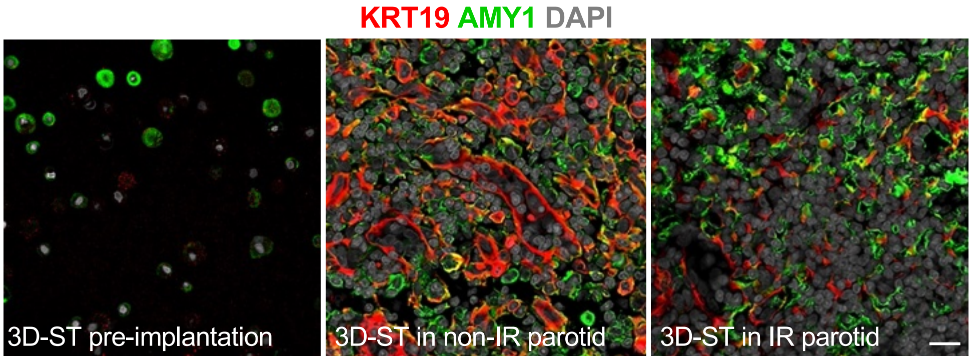
